# ReCap enables deep, copy number-scaled cysteine redox proteomics with minimal exogenous oxidation

**DOI:** 10.64898/2026.06.16.731536

**Authors:** James N. Cobley, Hao Jiang, Julia S. El-Sayed Moustafa, Melpomeni Platani, Zixin Kang, Beata Struckova, Russell Petty, Gillian P. Bates, Kerrin S. Small, Angus I. Lamond

## Abstract

Cysteine oxidation analyses require the preservation of the redox state present at harvest and quantitative scaling to relate oxidation to protein copy numbers, rather than only providing fractional oxidation data. Here, we present ‘ReCap’, a Redox Capture workflow combining Oxi-DIA, an enrichment-free isotope-encoded DIA workflow, with Oxi-Stop, a simple oxygen-exclusion strategy for cryopreserved tissue. In mouse brains, Oxi-DIA quantified 17,809 cysteine sites belonging to 6,085 protein groups in every sample, enabling matched measurements of residue-resolved oxidation and protein abundance. Atmospheric oxygen exposure during 14 days of cryopreservation distorted the measured cysteine redox state. The resultant increase of an estimated 5.3176 × 10^11^ *µg*^−1^ oxidised cysteine molecules was mitigated by Oxi-Stop, which minimised exogenous oxidation during cryopreservation. Copy-number scaling altered the interpretation of cysteine oxidation values. Although cysteine oxidation was detected across 2,371 sites and 1,439 proteins, 20 sites on abundant proteins accounted for 44% of the oxidised signal. ReCap advances redox proteomics from providing a site catalogue into a biologically weighted map of redox information, revealing cysteine oxidation as a sparse, ordered and quantitatively concentrated signal.

## Introduction

The ability of cysteine residues to modulate protein structure and function by serving as active sites, binding sites, or disulfide bonds depends on the oxidation state of the sulfur atom [1]. For example, metal binding in zinc-finger motifs largely depends on specific oxidation (−2) and protonation (i.e., thiolate) states. As a result, changing the oxidation of state the cysteine sulfur atom via electron exchange with suitable reactants like reactive oxygen species (ROS) can regulate protein structure and function [2–4].

Protein redox regulation is currently measured by enrichment-based proteomic approaches that quantify oxidation occupancy in unweighted dimensionless percentages [5–7]. Unweighted dimensionless percentages cannot, however, define the quantitative basis of redox regulation without copy number scaling [8]. Copy number scaling can profoundly alter the biological interpretation of the data [9,10]. For example, a 5% increase in oxidation on a cysteine residue in a low copy number protein that occurred despite the presence of orders-of-magnitude more abundant target sites implies specific oxidation mechanisms [11]. Copy number scale oxidation data are currently lacking because the enrichment steps needed for deep (≥ 10,000 *sites*) cysteine coverage discard the requisite protein group abundance information [12–14].

Regardless of the quantitative framework applied, any measurement of the cysteine redox state is only as reliable as the quality of the sample from which is it is derived [15]. This may be compromised by exogenous cysteine oxidation during sample cryopreservation and downstream processing. Exposure of samples to atmospheric oxygen ( ≈ 21% oxygen) may induce cysteine oxidation via transition metal ion-catalysed reactions that yield ROS [16] capable of oxidising cysteines [17]. Consequently [18,19], the measured redox state may differ substantially from the cysteine redox state that existed when the sample was harvested (*t*_*harvest*_ ≠ *t*_*analysis*_ ), confounding even technically rigorous experiments.

Here, we present redox capture (ReCap), a workflow comprising Oxi-DIA and Oxi-Stop. Oxi-DIA is an isotopologue-encoded data-independent acquisition (DIA) proteomic method for simultaneously quantifying residue-resolved cysteine redox states and protein groups. In mouse brains, Oxi-DIA provided deep quantitative copy number scaled measurement of cysteine residues (≥ 17,000 sites) and proteins (≈10,000 protein groups) without enrichment. Atmospheric oxygen exposure during cryopreservation confounded redox proteomics, spuriously amplifying the difference global difference between the left and right hemisphere by one order of magnitude. Oxi-Stop, wrapping a sample tightly in aluminium foil, minimised this redox drift by physically constraining oxygen diffusion during cryopreservation. ReCap provided the first copy number-scaled cysteine redox proteome data, with quantified oxidation shown as a sparse, structurally ordered signal.

## Results

### Oxi-DIA is an accurate and reproducible quantitative redox proteomic method

To quantify cysteine redox states with residue resolution, we developed Oxi-DIA (**Fig. 1**). In Oxi-DIA, reduced cysteines are first covalently alkylated with ‘light’ *N*-ethylmaleimide (NEM_L) at the point of tissue lysis. Leveraging established methods [20], reversibly oxidised cysteines, which were protected from alkylation at this step, are reduced using Tris-2-(carboxyethyl)phosphine (TCEP) and then alkylated with ‘heavy’ isotopologue-substituted *N*-ethyl-d_5_-maleimide (NEM_H). The 5-Dalton mass shift between NEM_L and NEM_H enables independent identification and quantification of the levels of the respective reduced, or reversibly oxidised, redox states for all detected cysteine residues. For each residue, the fractional oxidation level is calculated from the ratio of NEM_H-labelled signal to total NEM-labelled signal (NEM_L + NEM_H), yielding a bounded, dimensionless oxidation percentage. As this calculation is performed within protein groups that are quantified in parallel, the residue-level oxidation fraction is internally normalised and independent of variation in total protein abundances [21]. It can then subsequently be weighted and scaled using the protein group data acquired in parallel. We implemented Oxi-DIA using a single-shot, narrow-window 2-Thomson DIA [22] method on the Orbitrap-Astral™ MS instrument at a rate of 42-min per injection (i.e., throughput = 30 samples per day).

**Figure 1.**
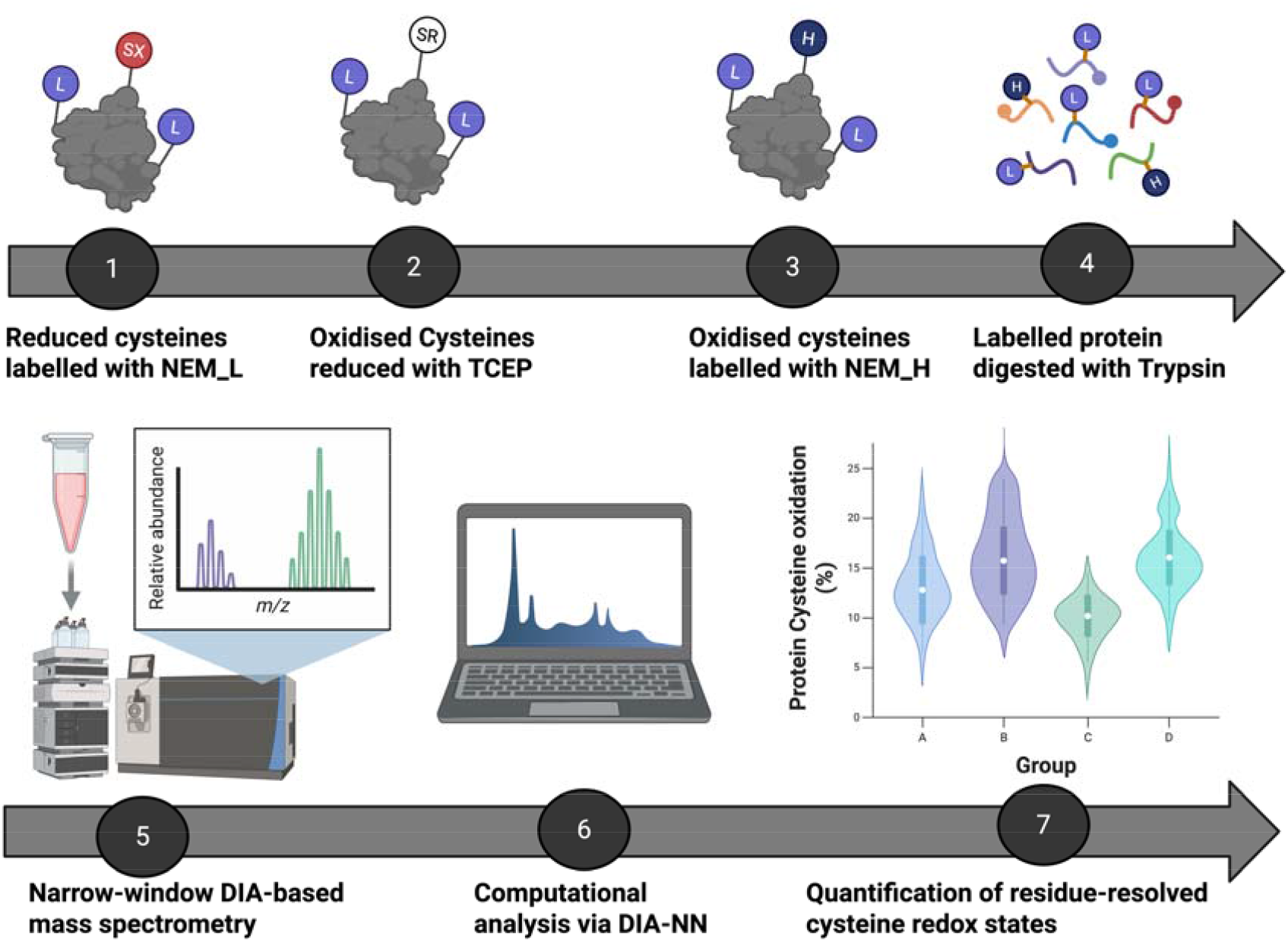
Overview of Oxi-DIA. 1. Reduced cysteines are alkylated with NEM_L at lysis via a covalent thioether bond. 2. After removing NEM_L (omitted for clarity), reversibly oxidised cysteines are chemically reduced using Tris-2(carboxyethyl)phosphine (TCEP). 3. These newly reduced cysteines are then labelled with NEM_H (+ 5 Daltons) to isotopically encode the difference between the reduced and oxidised state. 4. Labelled proteins are then digested at lysine and arginine residues via Trypsin + Lys-C. 5. The resultant peptides are analysed via narrow-window DIA. 6. The raw files are analysed using DIA-NN. 7. Open-sourced scripts are used to analyse the DIA-NN output files, such as to calculate mean cysteine oxidation.

To validate Oxi-DIA, we generated an eight-point standard curve spanning 0–100% reduced cysteine, by mixing equal amounts of fully NEM_H-labelled and fully NEM_L-labelled mouse brain lysates at defined ratios. Mouse brain lysates were used to support future studies on neurodegeneration [23,24]. After Oxi-DIA-based data acquisition, raw files were analysed using DIA-NN [25]. Oxi-DIA demonstrated excellent quantitative accuracy across the full dynamic range (**Fig. 2**).

**Figure 2.**
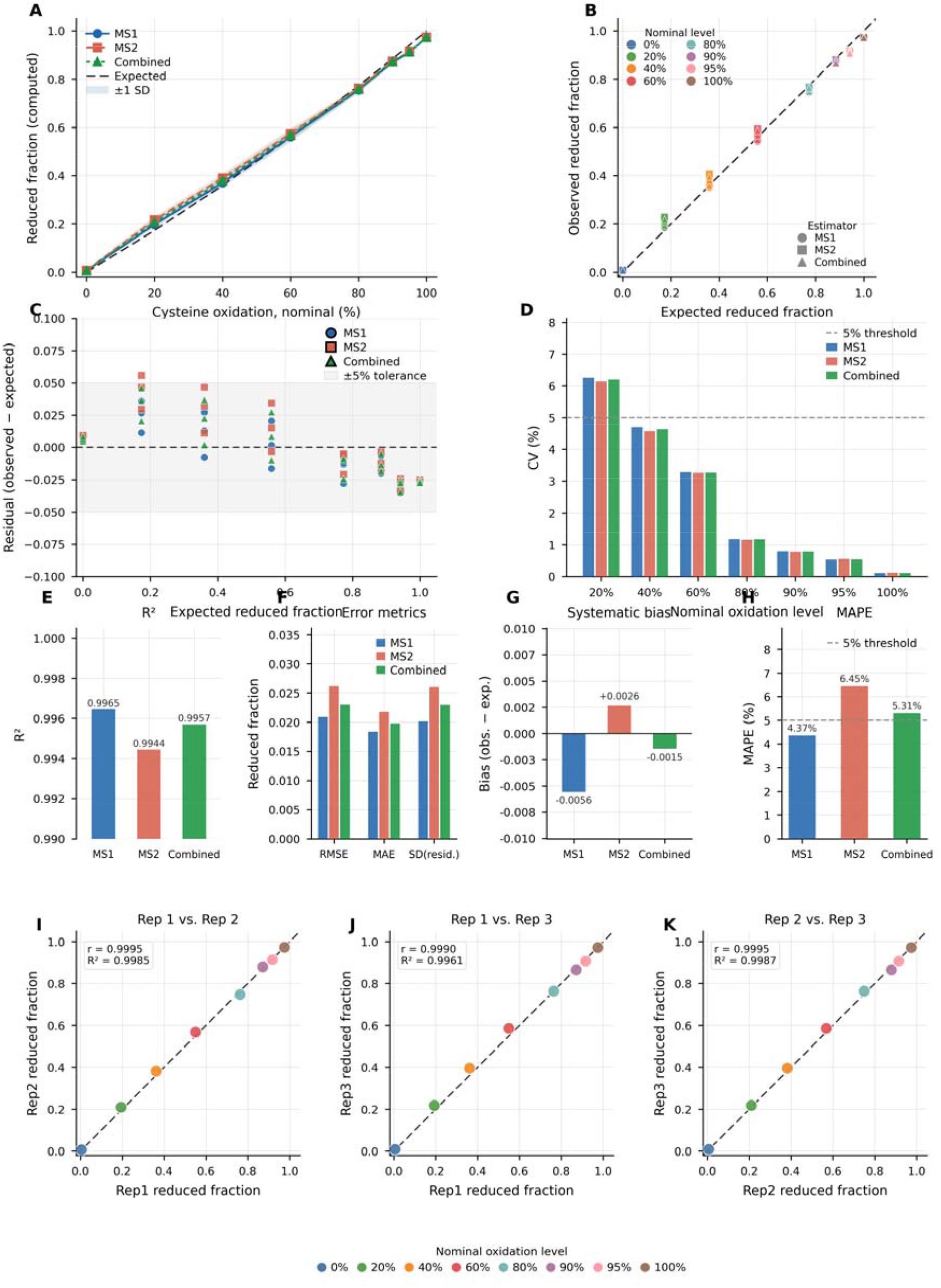
Validation of Oxi-DIA for quantitative cysteine redox proteomics across a defined mixing series. A defined mixture series of isotopically labelled (heavy, fully oxidised) and unlabelled (light, fully reduced) cysteine-containing peptides was prepared at nominal oxidation levels spanning 0–100%, with three technical replicates per level. Reduced fractions were computed using three estimators: MS^1^ (light peptide counts relative to all detected cysteine peptides), MS^2^ (light relative to labelled peptides only), and a Combined average of both. **(A)** Mean reduced fraction computed by each estimator plotted against the nominal cysteine oxidation level. Shaded bands represent ±1 standard deviation across technical replicates (*n* = 3). The dashed line indicates the model-expected reduced fraction, adjusted for the measured unlabelled peptide background estimated from the 0% and 100% reference conditions. **(B)** Per-replicate observed reduced fraction plotted against the model-expected value for each estimator (circle, MS^1^; square, MS^2^; triangle, Combined). Points are coloured by nominal oxidation level. Proximity to the identity line (dashed) indicates quantitative accuracy across the full dynamic range. **(C)** Residuals (observed minus expected reduced fraction) plotted against the expected value for each estimator. The shaded band denotes the ±5% tolerance interval. All data points fall within or close to this band, indicating no systematic non-linearity across the quantification range. **(D)** Coefficient of variation (CV%) per nominal oxidation level for each estimator, excluding the 0% condition (where CV is uninformative due to a near-zero denominator). The dashed line marks the 5% CV threshold. CV falls below 5% for all levels ≥40%. **(E)** Coefficient of determination (R^2^) for each estimator relative to the expected reduced fraction. Axis is truncated to 0.990–1.000 to resolve inter-estimator differences; all estimators achieve R^2^ > 0.994. **(E)** Absolute error metrics — root mean square error (RMSE), mean absolute error (MAE), and standard deviation of residuals SD(resid.) — for each estimator. All values are <0.027 reduced fraction units across the full range. **(G)** Systematic bias (mean residual) per estimator. MS1 exhibits a small negative bias (−0.006), MS^2^ a small positive bias (+0.003), and the Combined estimator achieves near-zero bias (−0.001), supporting its use as the primary quantitative readout. **(H)** Mean absolute percentage error (MAPE) per estimator. The dashed line marks the 5% threshold. MS^1^ achieves the lowest MAPE (4.37%), with Combined at 5.31%. **(I–K)** Pairwise technical replicate correlations for the Combined estimator across all oxidation levels. Pearson correlation coefficient (*r*) and coefficient of determination (R^2^) are annotated per panel. All replicate pairs achieve r ≥ 0.999 and R^2^ ≥ 0.996, demonstrating exceptional analytical reproducibility. Points are coloured by nominal oxidation level as indicated in the shared legend.

Observed reduced fractions tracked closely with model-expected values for all three estimators, yielding R^2^ values of 0.9965 (MS^1^), 0.9944 (MS^2^), and 0.9957 (Combined) (**Fig. 2E**). Absolute error was low across all estimators, with root mean square error (RMSE) of 0.021 (MS^1^), 0.026 (MS^2^), and 0.023 (Combined), and mean absolute error (MAE) of 0.018, 0.022, and 0.020, respectively (**Fig. 2F**). Residuals were tightly distributed around zero across the quantification range, with all individual measurements falling within or close to the ±5% tolerance interval (**Fig. 2C**). Systematic bias was negligible for all estimators (MS^1^: −0.006; MS^2^: +0.003; Combined: −0.001), with the Combined estimator achieving near-zero bias through partial cancellation of opposing MS^1^ and MS^2^ directional errors (**Fig. 2G**). Mean absolute percentage error was 4.37% (MS^1^), 6.45% (MS^2^), and 5.31% (Combined) (**Fig. 2H**). Inter-replicate precision was high across all nominal oxidation levels, with CV% falling below 5% for all levels at or above 40% reduced (**Fig. 2D**). Technical reproducibility was exceptional, with all pairwise replicate correlations achieving r ≥ 0.999 and R^2^ ≥ 0.996 across the full range (**Fig. 2I–K**). In parallel, Oxi-DIA quantified a mean of 10,562 unique protein groups per technical replicate, with high-quality spectral data maintained across the acquisition series [26] (**Supplementary Figure 1**).

### Oxi-DIA achieved deep, multidimensional cysteine redox proteomics

Next, we performance benchmarked and externally validated Oxi-DIA in freshly lysed HEK293 lysates. We first validated the minimisation of endogenous lysis-induced oxidation by using chelators to limit transition metal ion catalysed autoxidation [17] (**Supplementary Figure 2**). Omitting sample sonication and boiling minimised cysteine oxidation, likely by preventing acoustic cavitation mediated hydroxyl radical generation [27] and the homolysis of disulfide bonds to form sulfur radicals [28], respectively (**Supplementary Figure 2**). After the NEM_H labelling step, where reoxidation is minimised by TCEP and chelators, cysteine oxidation during the rest of the proteomic workflow is improbable owing to alkylation.

Oxi-DIA generated reproducibly deep cysteine redox proteome coverage across all six HEK293 samples (**Fig. 3**). Across the samples, cysteine redox-state measurements were dominated by fully or near-fully reduced sites, with an overall mean oxidation of 0.91% and a median oxidation of 0.00%, indicating that reversible cysteine oxidation was sparse relative to the measured cysteine proteome (**Fig. 3A-B**). Oxi-DIA consistently quantified >22,000 cysteine sites per sample (range: 22,451-23,592, **Fig. 3C**). These sites mapped to a highly reproducible cysteine-containing protein space, with 6,287–6,502 unique protein groups containing detected cysteines per sample (**Fig. 3D**). Cysteine labelling was near-complete, with precursor-level labelling efficiencies of 99.0–99.8% (**Fig. 3E**). Total proteome depth was also stable across samples, with approx. 10,000 protein groups detected per sample, and 165,360–170,419 unique stripped peptide sequences detected per sample (**Fig. 3F-G**). Hence, Oxi-DIA achieved deep multidimensional analysis of cysteines (approx. 20,000) and protein groups (approx. 10,000) without enrichment.

**Figure 3.**
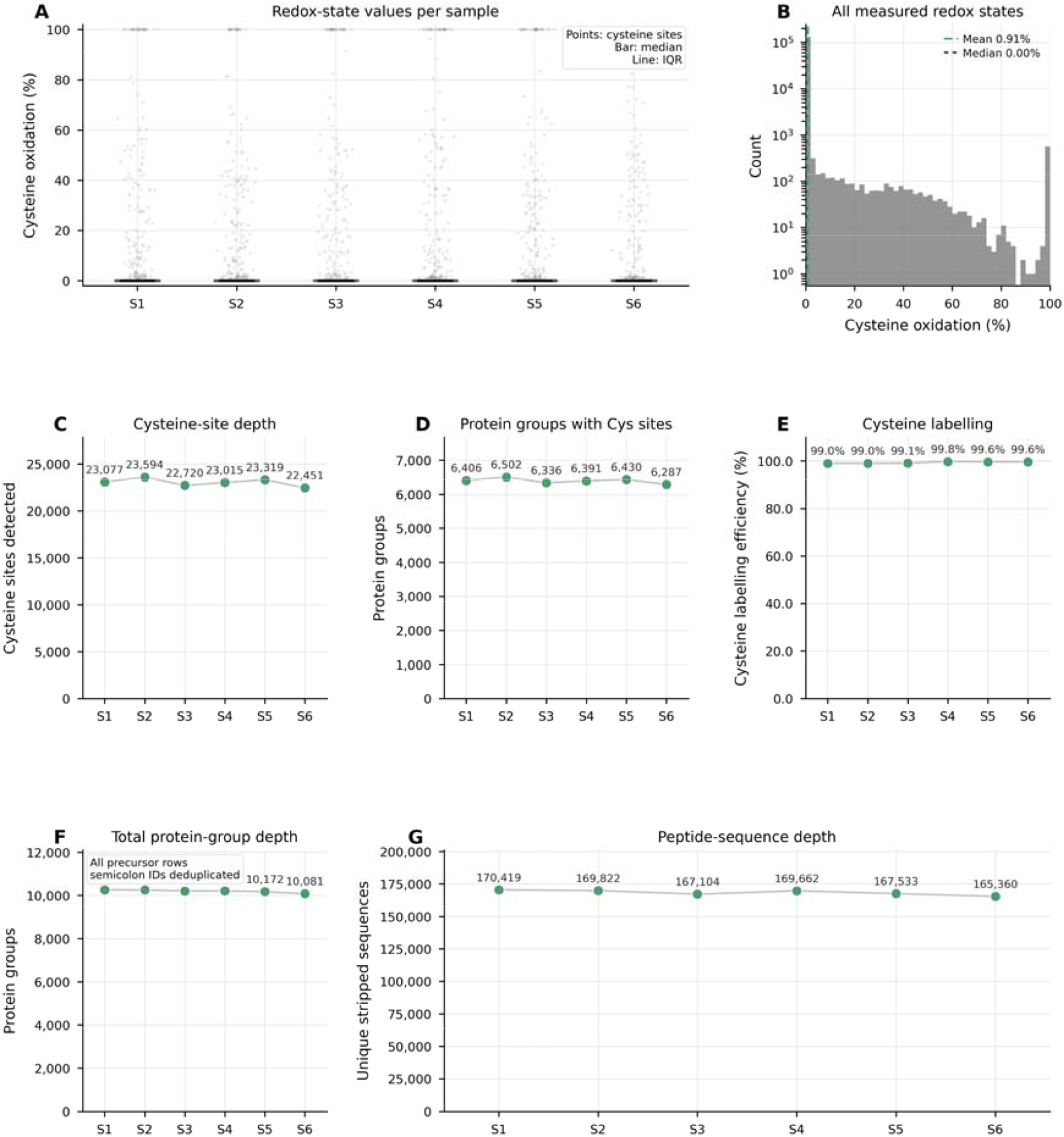
Sample-level depth, labelling efficiency, and redox-state structure of the ReCap dataset. **(A)** Distribution of cysteine redox-state values for each sample. Each point represents a measured cysteine-site oxidation value; horizontal bars indicate the sample median and vertical lines indicate the interquartile range. Across all samples, most cysteine sites were fully or near-fully reduced, with a sparse subset of oxidised sites extending across the full 0–100% redox-state range. **(B)** Histogram of all measured cysteine redox-state values across the dataset, showing a strongly reduced global distribution with a mean oxidation of 0.91% and a median oxidation of 0.00%. **(C)** Cysteine-site depth per sample, showing consistently deep redox coverage across all six samples, with approximately 22,451–23,594 cysteine sites detected per sample. **(D)** Number of unique protein groups containing detected cysteine redox sites per sample, ranging from 6,287 to 6,502 protein groups. **(E)** Precursor-level cysteine labelling efficiency per sample, calculated as the percentage of cysteine-containing peptide precursors labelled with either NEM_L or NEM_H. Labelling efficiency was near-complete across samples, ranging from 99.0% to 99.8%. **(F)** Total protein-group depth per sample calculated from all detected precursor rows, with semicolon-delimited protein identifiers deduplicated. Approximately 10,000 protein groups were detected per sample. **(G)** Unique stripped peptide-sequence depth per sample, showing 165,360–170,419 unique stripped sequences per sample.

Chromatographically, 1,038,988 non-cysteine peptide precursors, 155,184 NEM_L-labelled reduced cysteine precursors, and 4,451 NEM_H-labelled reversibly oxidised cysteine precursors were detected across the chromatographic gradient (**Fig. 4A**). The lower number of NEM_H-labelled precursors was therefore consistent with a sparse reversibly oxidised cysteine pool rather than a general failure of peptide detection. As expected, matched NEM_L/NEM_H forms of the same peptide showed near-identical chromatographic behaviour, with 3,833 paired precursors lying close to the identity line for retention time correlation (*r* = 0.998), and a small mean retention time difference of −0.068 min (**Fig. 4B-C**). Isotope encoding was also preserved at the mass level: 3,748 of 3,833 matched pairs fell within the expected mass-shift window, with an observed median mass difference of 5.029 Da compared with the theoretical 5.031 Da (**Fig. 4D**). The NEM_H signal occupied a broad MS1 intensity range spanning 4.2 orders of magnitude, compared with 5.6 orders of magnitude for NEM_L. At the NEM_H limit-of-detection threshold, 99% of both NEM_H and NEM_L precursors remained detectable, and 100% of NEM_H precursors fell within the measured NEM_L intensity range (**Fig. 4E-F**). Hence, the relative scarcity of NEM_H-labelled peptides was not explained by a technical limit-of-detection artefact. Instead, the data support the interpretation that Oxi-DIA resolved a genuinely low-abundance oxidised cysteine sub-proteome within the same chromatographic and quantitative intensity space as the reduced cysteine proteome.

**Figure 4.**
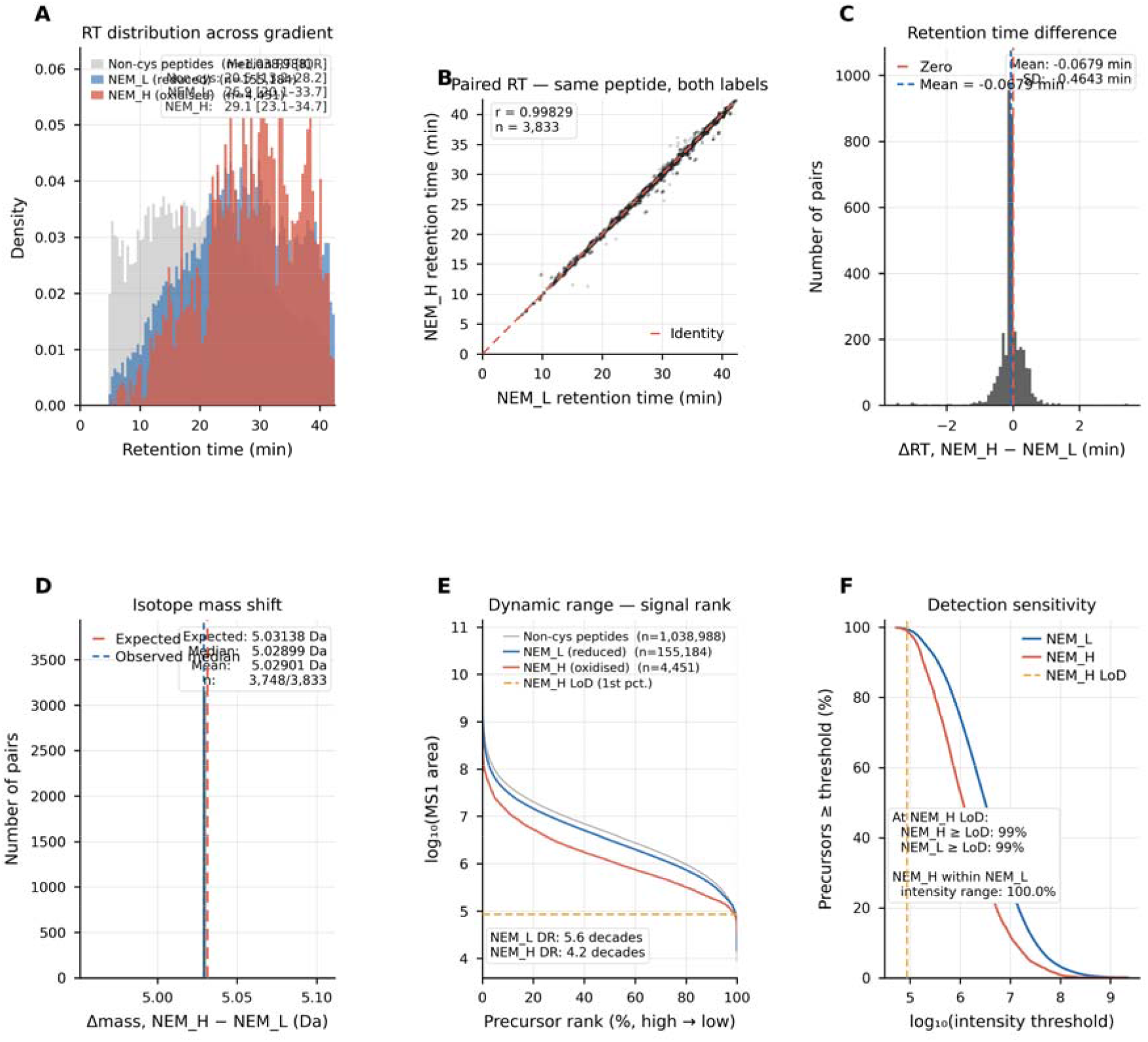
LC–MS validation of isotope-encoded cysteine redox proteomics. **(A)** Retention-time distributions of non-cysteine peptides, NEM_L-labelled reduced cysteine peptides, and NEM_H-labelled reversibly oxidised cysteine peptides across the LC gradient. All peptide classes were distributed across the chromatographic run. **(B)** Paired retention-time comparison for matched NEM_L/NEM_H forms of the same peptide, showing near-identity co-elution across 3,833 matched precursor pairs. **(C)** Distribution of paired retention-time differences, calculated as NEM_H − NEM_L, centred close to zero with a mean offset of −0.0679 min. **(D)** Paired precursor mass differences were concentrated around the expected isotope mass shift for NEM_H relative to NEM_L. Within the plotted mass-shift window, 3,748 of 3,833 matched pairs were retained, with a median observed shift of 5.0289 Da compared with the theoretical shift of 5.0314 Da. **(E)** Signal-rank analysis of MS1 precursor intensities showed that NEM_H-labelled oxidised cysteine peptides occupied a broad quantitative range, spanning 4.2 orders of magnitude, compared with 5.6 orders of magnitude for NEM_L-labelled reduced cysteine peptides. **(F)** Detection-sensitivity analysis showed that, at the NEM_H limit-of-detection threshold, 99% of both NEM_H and NEM_L precursors remained above threshold, and 100% of NEM_H precursors fell within the measured NEM_L intensity range.

After benchmarking Oxi-DIA, we used two complementary approaches to externally validate the method (**Supplementary Figure 3**). First, we covalently captured reversibly oxidised cysteines, before digesting the enriched (i.e., oxiforms bearing at least one reversibly oxidised residue) and eluent (i.e., reduced) fractions. Redox state-dependent fractionation was confirmed by background fluorescent maleimide (F-MAL) labelling in the presence of TCEP in the eluent and substantial F-MAL labelling in the presence of TCEP in the enriched fraction (**Supplementary Figure 3**). The 0.91% cysteine redox state was qualitatively recapitulated offline by measuring the peptide yield in each fraction, which was substantially higher in the eluent (i.e., reduced fraction). Second, we quantitatively compared Oxi-DIA against an orthogonal microplate assay, RedoxiFluor [29]. In RedoxiFluor, the NEM_L and NEM_H labels were replaced with spectrally distinct fluorophores. We observed near-perfect correspondence between Oxi-DIA and RedoxiFluor in HEK293 cells (**Supplementary Figure 3**). Orthogonal validation excluded any quantitative measurement issue intrinsic to Oxi-DIA [30].

### Oxi-Stop: A simple oxygen-exclusion strategy

Irrespective of whether Oxi-DIA or any other method is applied, quantitative redox measurements of a tissue can be compromised before the sample was even lysed due to atmospheric oxygen-induced exogenous oxidation during cryopreservation [31]. Given the practical inability to hermetically seal samples, we developed Oxi-Stop, a simple aluminium foil-based oxygen exclusion strategy. Aluminium foil is widely used to preserve food by limiting oxidation: its chemically inert aluminium oxide surface forms a dense, nonporous barrier that is effectively impermeable to oxygen at thicknesses ≥15 µm [32].

In support of this interpretation, we used a model oxidised thiol 5,5’dithiobis-(2-nitrobenzoic acid) (DTNB [33]) to determine if aluminium foil could act as a reducing agent. Upon reduction, DTNB forms the colorimetric product 2-nitro-5-thiobenzoic acid (TNB). The level of TNB formation when DTNB was incubated with aluminium foil in either the solid, or aqueous state, was identical to the negative control material: commercially passivated plastic (**Supplementary Figure 3**). In contrast, TCEP reduced DTNB to TNB. Hence, we concluded that Oxi-Stop, oxygen exclusion by aluminium foil wrapping, can physically constrain atmospheric oxygen diffusion during cryopreservation.

### Exogenous oxidation during cryopreservation spuriously amplified a biological difference

Having established that aluminium foil likely provides an inert physical barrier to oxygen diffusion, we isolated atmospheric oxygen exposure as a preanalytical variable in tissue redox proteomics. The key experimental requirement was to devise a paired system with a plausible biological redox asymmetry that could then be manipulated by the storage condition. Accordingly, we asked whether the cysteine proteome measurably between the left and right brain hemispheres of the same mouse to define if redox regulation displays sidedness [34]. Using hemispheres that were both stored identically using Oxi-Stop, we observed a small difference in global cysteine oxidation between hemispheres (Δ = 0.04%; **Supplementary Figure 3**). This established the paired-hemisphere system as a model for appraising exogenous oxidation as the spurious amplification of Δ = 0.04% difference between hemispheres.

Freshly isolated mouse brain right hemispheres were tightly wrapped in aluminium foil (Oxi-Stop), prior to placement in a cryovial. The left hemispheres were placed directly into a cryovial, exposed to atmospheric oxygen during cryopreservation (Control, CTRL). All samples were then flash-frozen and stored at −70°C for 14 days before identical Oxi-DIA processing and analysis. In this experiment, Oxi-DIA quantified 17,809 cysteine sites across 6,085 protein groups in every sample (i.e., no missing values). The mean cysteine coverage for these protein groups (detected vs. total cysteines) was 30.3% (**Supplementary Figure 4**).

If the difference in storage conditions in this experiment did not significantly affect measured redox values, then a difference in global cysteine redox level of 0.04% between the left and right brain hemisphere samples would be expected. Instead, the cysteine redox state of the brain hemispheres wrapped in aluminium foil was 2.68%-oxidised, compared with 2.53%-oxidised in the CTRL group (**Fig. 5A**). The resultant redox state change (Δ = −0.14%) exceeded the previously measured biological redox asymmetry between the hemispheres (**Fig. 5B**). Hence, storage spuriously amplified the biological difference between the hemispheres by an order of magnitude, confounding redox proteomics.

**Figure 5.**
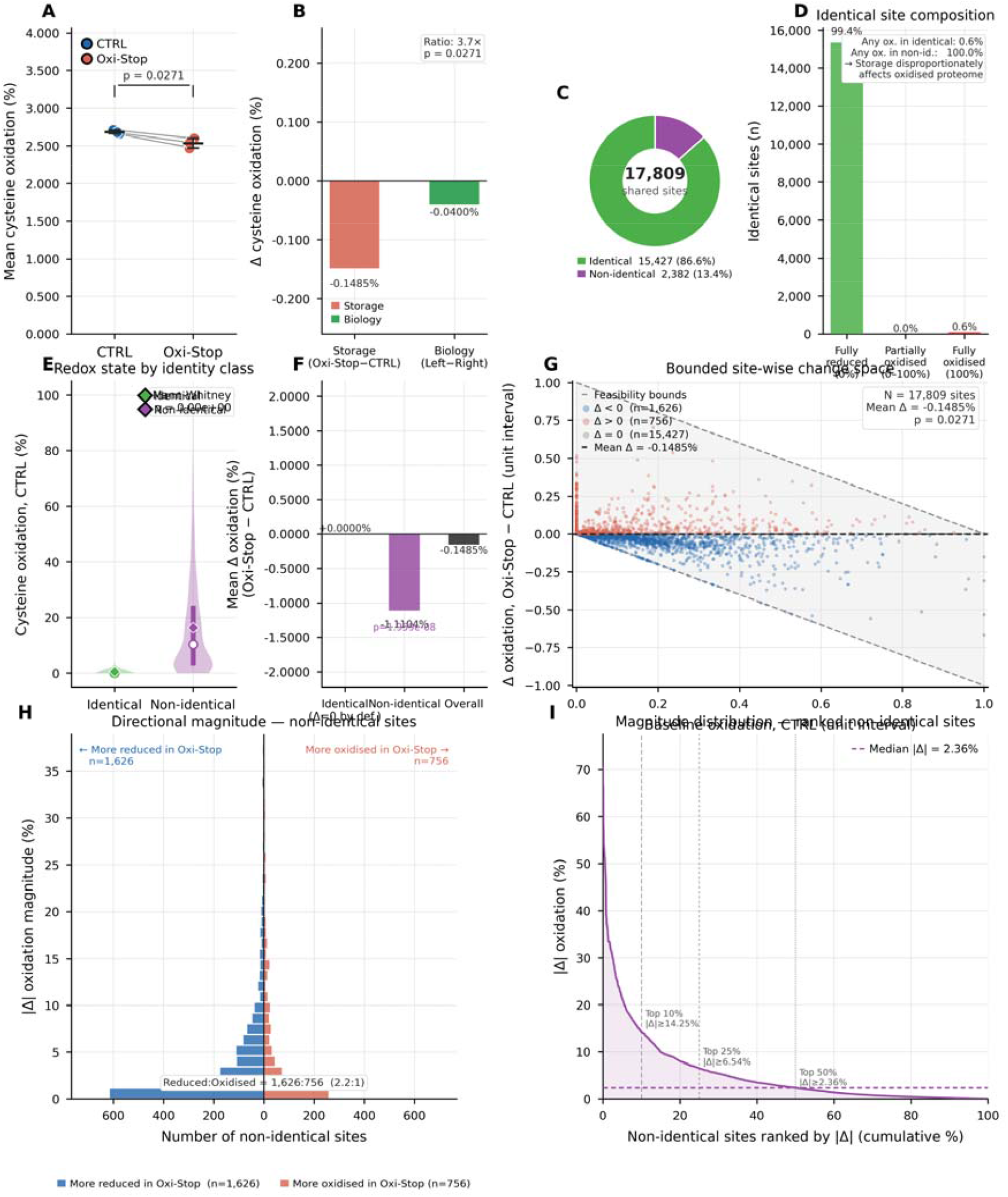
Site-level cysteine redox proteome analysis of Oxi-Stop storage versus physiological control conditions. A defined cohort of 17,809 cysteine sites detected across all 18 injections (3 biological replicates × 3 technical injections per condition) was used for all analyses. Technical replicates were averaged within each biological replicate prior to statistical testing; all comparisons use n = 3 paired biological means as the unit of analysis. **(A)** Mean cysteine oxidation per biological replicate for CTRL and Oxi-Stop conditions. Lines connect paired biological replicates. Error bars represent ± 1 SD. Paired t-test: p = 0.0271. **(B)** Comparison of the storage-induced delta in mean cysteine oxidation (Oxi-Stop − CTRL, −0.149%) against the hemispheric biological delta (Left − Right, −0.040%). The storage effect is 3.7× the magnitude of the endogenous hemispheric biological variation, confirming that Oxi-Stop induces a small but reproducible and measurable change in the cysteine redox proteome. **(C)** Proportion of shared cysteine sites classified as identical (Δ = 0.000%, n = 15,427, 86.6%) or non-identical (Δ ≠ 0%, n = 2,382, 13.4%) between conditions. **(D)** Composition of identical sites by oxidation state: 99.4% are fully reduced (0%), 0.6% are partially oxidised (0–100%), and 0.0% are fully oxidised (100%). Inset annotation indicates that 100% of non-identical sites carry any detectable oxidation in either condition versus 0.6% of identical sites, demonstrating that storage-induced changes are concentrated within the oxidised fraction of the cysteine proteome. **(E)** Distribution of cysteine oxidation levels at baseline (CTRL) for identical and non-identical sites shown as violin plots with interquartile range and median. Diamond markers indicate group means. Non-identical sites exhibit significantly higher baseline oxidation than identical sites (Mann-Whitney U test, p reported in panel), consistent with storage preferentially perturbing cysteines that carry detectable oxidation under physiological conditions. **(F)** Mean delta (Oxi-Stop − CTRL) stratified by site identity class. Identical sites have a mean delta of 0.000% by formal construction. Non-identical sites show a mean delta of −1.104%, and the overall mean delta across all shared sites is −0.149%. **(G)** Bounded site-wise change space. Each point represents one cysteine site plotted as baseline CTRL oxidation (x-axis, unit interval) against the change in oxidation upon storage (y-axis, unit interval). The feasible region (grey shading) is bounded above by 1 − x and below by −x, reflecting the physical constraint that oxidation cannot exceed 100% or fall below 0%. Points are coloured by direction of change: blue, Δ < 0 (more reduced in Oxi-Stop, n = 1,626); red, Δ > 0 (more oxidised in Oxi-Stop, n = 756); grey, Δ = 0 (identical, n = 15,427). The dashed horizontal line indicates the mean Δ = −0.149%. **(H)** Directional magnitude histogram for non-identical sites. The y-axis represents the absolute magnitude of the oxidation change; bars extend left (blue) for sites more reduced in Oxi-Stop and right (red) for sites more oxidised. Most non-identical sites are more reduced under Oxi-Stop storage conditions per the negative overall mean delta. **(I)** Magnitude distribution of non-identical sites ranked in descending order of |Δ|. Reference lines indicate the minimum |Δ| required to fall within the top 10% (≥14.25%), 25% (≥6.54%), and 50% (≥2.36%, median) of non-identical sites by effect size. The rapid decay indicates that the storage effect is dominated by a small number of sites with large oxidation changes, while most nonidentical sites exhibit modest shifts.

### Storage disproportionately distorted the oxidised cysteine sub-proteome

The global shift in mean oxidation between the groups concealed a much stronger site-level effect. We therefore classified each cysteine as either identical, where oxidation was unchanged between Oxi-Stop and CTRL (Δ = 0%), or non-identical, where oxidation differed between conditions (Δ ≠ 0%). This partition separated the cysteine proteome into two distinct regimes [35].

Of the 17,809 cysteine sites quantified in every sample, 15,427 sites were identical between Oxi-Stop and CTRL, corresponding to 86.6% of the measured cysteine proteome (**Fig. 5C**). Among the identical cysteines, 99.4% were 0%-oxidised in both conditions, whereas only 0.6% showed any oxidation, and these were almost entirely fully oxidised sites (**Fig. 5D**). No partially oxidised site was identical. Even a seemingly adventitious process—exogenous oxidation—did not act randomly across the proteome, its effect was concentrated into a small number of sites.

In contrast, the oxidised sub-proteome was disproportionately unstable. The remaining 2,382 cysteine sites, representing 13.4% of all quantified sites, were non-identical between Oxi-Stop and CTRL (**Fig. 5C**). Unlike the identical population, these sites occupied the redox-active state space: they showed measurable oxidation in at least one condition and accounted for essentially all partially oxidised cysteines. Consequently, the difference between Oxi-Stop and CTRL was concentrated within the subset of sites most likely to be interpreted as biologically redox regulated [36] (**Fig. 5E**).

This partition also explained why the global mean underestimated the magnitude of the storage effect. Identical sites, which were overwhelmingly fully reduced, had a mean Δ of 0%. By contrast, non-identical sites showed a much larger mean shift, while the overall proteome-wide mean was diluted by the large invariant reduced population (**Fig. 5F**).

The direction of change further supported an oxygen-exposure effect. Across all sites, 1,626 cysteines were more reduced in Oxi-Stop than in CTRL, whereas 756 cysteines were more oxidised in Oxi-Stop than in CTRL (**Fig. 5G-H**). Thus, sites showing lower oxidation under Oxi-Stop outnumbered sites showing higher oxidation by approximately 2.1:1. The dominant direction therefore corresponded to increased oxidation in the oxygen-exposed CTRL samples. The bounded analysis also showed that these changes occurred within the mathematically constrained interval imposed by the initial oxidation state: sites close to 0% could only gain oxidation, whereas sites closer to 100% had progressively less oxidative headroom (**Fig. 5G**).

Moreover, the magnitude of the non-identical changes was highly uneven. Ranking non-identical cysteines by absolute oxidation difference revealed a long-tailed distribution, with a median |Δ| of 2.36%, the top 25% of sites changing by at least 6.54%, and the top 10% changing by at least 14.25% (**Fig. 5I**).

### The exogenous storage signature is superimposed on the endogenous membrane-interface redox landscape

After mapping all cysteines in the mouse reference proteome to UniProt positional annotations (**Supplementary Figure 4**), we asked whether storage-sensitive cysteines were enriched within functionally annotated redox space. Of the cysteines quantified by Oxi-DIA, 24.3% of non-identical sites carried a functional annotation, compared with 10.2% of invariant cysteines, corresponding to a 2.84-fold enrichment (**Fig. 6A**). This enrichment was directionally asymmetric and concentrated in sites that were more oxidised in CTRL than in Oxi-Stop, consistent with protection by oxygen exclusion. Functionally annotated non-identical sites also showed larger storage-dependent redox displacement than unannotated non-identical sites: mean oxidation decreased from 21.6% in CTRL to 18.1% in Oxi-Stop, corresponding to a mean Δ of −3.43%, compared with −0.37% for unannotated non-identical cysteines (**Fig. 6B**). Feature-class analysis showed that this signal was driven primarily by disulfide-annotated cysteines, which were enriched 4.24-fold among non-identical sites (**Fig. 6C-D**). However, stratification by evidence level showed that this was not a canonical structural disulfide signature: PDB-supported disulfide annotations represented only a small and similar fraction of disulfide-annotated sites in identical and non-identical classes. Instead, the non-identical disulfide-annotated pool was enriched in predicted or sequence-model annotations, indicating that storage oxygen preferentially distorted a distinct disulfide-annotated redox-active pool rather than simply oxidising structurally resolved disulfide bonds.

**Figure 6.**
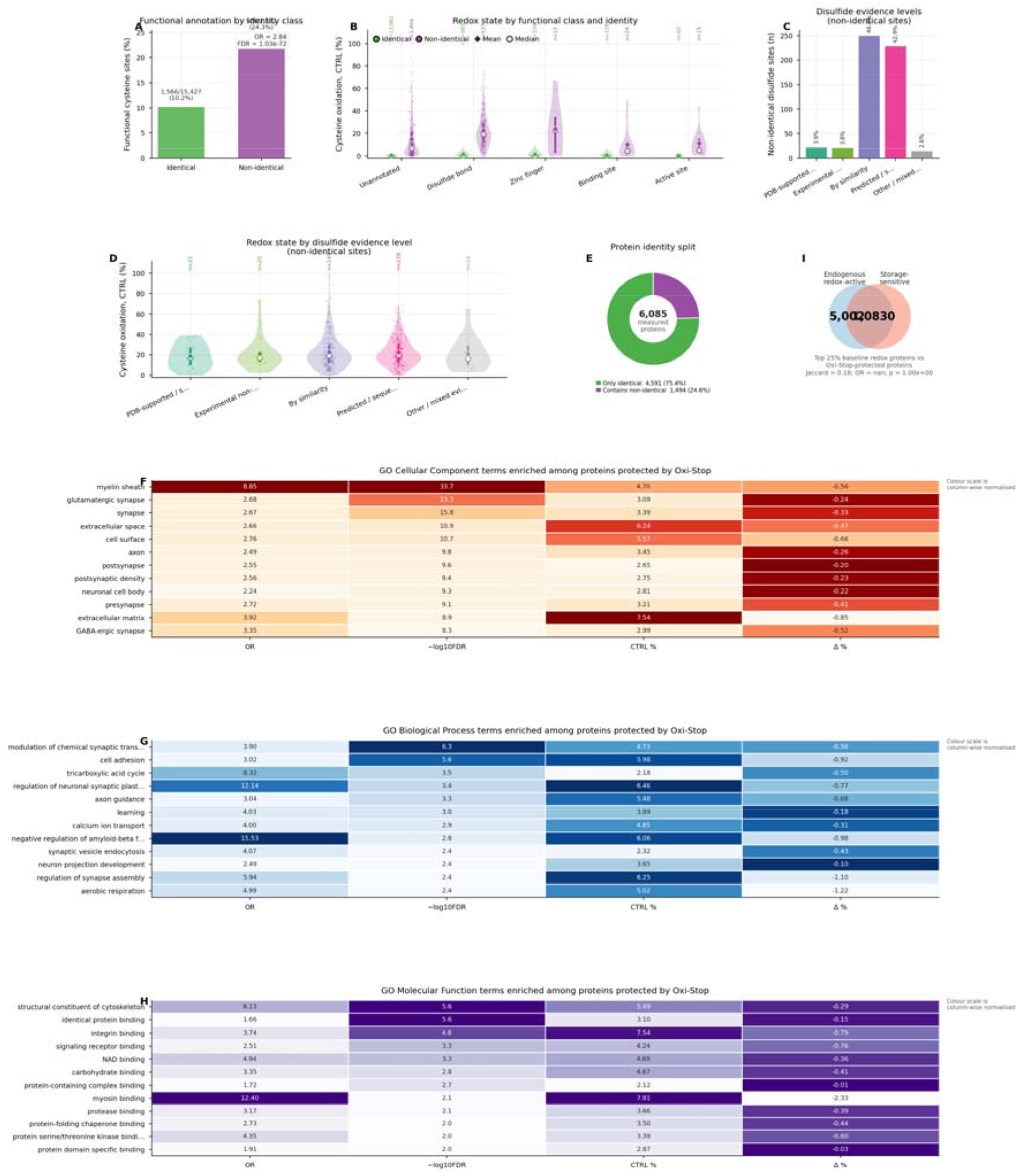
Exogenous storage-induced oxidation is superimposed on the endogenous membrane-interface redox signature. **(A)** Functional cysteine annotations were enriched among non-identical storage-sensitive cysteines. Bars show the percentage of identical and non-identical cysteine sites carrying at least one UniProt positional functional annotation. Non-identical cysteines were 2.84-fold enriched for functional annotation. **(B)** CTRL redox state distributions for cysteine classes split by identity class. Identical cysteines were concentrated at the fully reduced boundary, whereas non-identical cysteines occupied higher and more broadly distributed oxidation states across unannotated and functionally annotated classes. Diamonds indicate means and open circles indicate medians. **(C)** Evidence-level composition of disulfide-annotated non-identical cysteines. The non-identical disulfide-annotated pool was dominated by by-similarity and predicted/sequence-model annotations, whereas PDB-supported structural disulfides represented only a small fraction. **(D)** CTRL redox state distributions of non-identical disulfide-annotated cysteines stratified by evidence level. **(E)** Protein-level identity split showing the number of measured protein groups containing only identical cysteines or at least one non-identical cysteine. **(F)** GO Cellular Component enrichment heatmap for proteins containing cysteine sites protected by Oxi-Stop. Terms are shown with odds ratio, −log10(FDR), baseline CTRL oxidation, and mean storage-dependent Δ oxidation. Enriched terms were dominated by membrane-interface compartments including myelin sheath, synapse, extracellular space, cell surface, extracellular matrix, axon, dendrite, and plasma membrane. **(G)** GO Biological Process enrichment heatmap for the same protein set, highlighting neuronal, synaptic, amyloid-related, and respiratory processes. **(H)** GO Molecular Function enrichment heatmap showing enriched functional terms including cytoskeletal binding, integrin binding, myosin binding, NAD binding, and protein kinase binding. **(I)** Overlap between proteins with high baseline cysteine oxidation and proteins containing Oxi-Stop-protected cysteine sites, illustrating the superimposition of the endogenous redox-active protein space and the storage-sensitive protein space. Across panels F–H, negative Δ values indicate lower oxidation in Oxi-Stop than CTRL, consistent with protection from exogenous oxygen exposure during cryopreservation.

We next asked whether this storage-sensitive functional cysteine pool mapped onto coherent protein-level biology. Among 6,085 measured protein groups, 1,494 contained at least one non-identical cysteine (**Fig. 6E**). Proteins containing Oxi-Stop-protected sites were strongly enriched for membrane-interface cellular component terms, including myelin sheath, synapse, glutamatergic synapse, extracellular space, cell surface, postsynapse, presynapse, extracellular matrix, axon, dendrite, and plasma membrane (**Fig. 6F**). These same terms also showed elevated baseline cysteine oxidation and negative Δ values, indicating that they were both endogenously oxidised in CTRL and protected by Oxi-Stop. Biological process and molecular function enrichment extended this pattern to synaptic transmission, axon guidance, neuron projection development, amyloid-β regulation, aerobic respiration, integrin binding, cytoskeletal binding, myosin binding, NAD binding, and protein kinase binding (**Fig. 6G-H**). Hence, exogenous oxygen exposure did not generate an orthogonal technical artefact. It acted on the same membrane-, synapse-, myelin-, extracellular-, and cytoskeletal-associated redox landscape in which endogenous cysteine oxidation was already concentrated (**Fig. 6I**).

### Copy-number scaling resolved cysteine oxidation as a sparse signal

The apparent magnitude of the Oxi-Stop effect depended strongly on whether cysteine oxidation was interpreted as an unweighted site percentage or as a copy-number-scaled molecular signal. In the shared-site analysis, the unweighted mean suggested a small between-group shift of −0.1485 percentage points. However, after TPA-based copy-number scaling [37], the estimated cysteine redox state was 5.52% in CTRL and 4.95% in Oxi-Stop, corresponding to a larger shift of −0.575 percentage points (**Fig. 7A-B**). Thus, abundance scaling increased the apparent between-group difference by 3.9-fold. When converted into molecular units, this difference corresponded to 4.81 × 10^12^ oxidised cysteine molecules per µg protein in CTRL and 4.31 × 10^12^ in Oxi-Stop. Therefore, exclusion of atmospheric oxygen reduced the oxidised cysteine channel by 5.01 × 10^11^ molecules per µg protein, equivalent to a 10.4% decrease (**Fig. 7C**). A change that appeared modest on a dimensionless percentage scale therefore represented a substantial molecular displacement in the cysteine redox channel.

**Figure 7.**
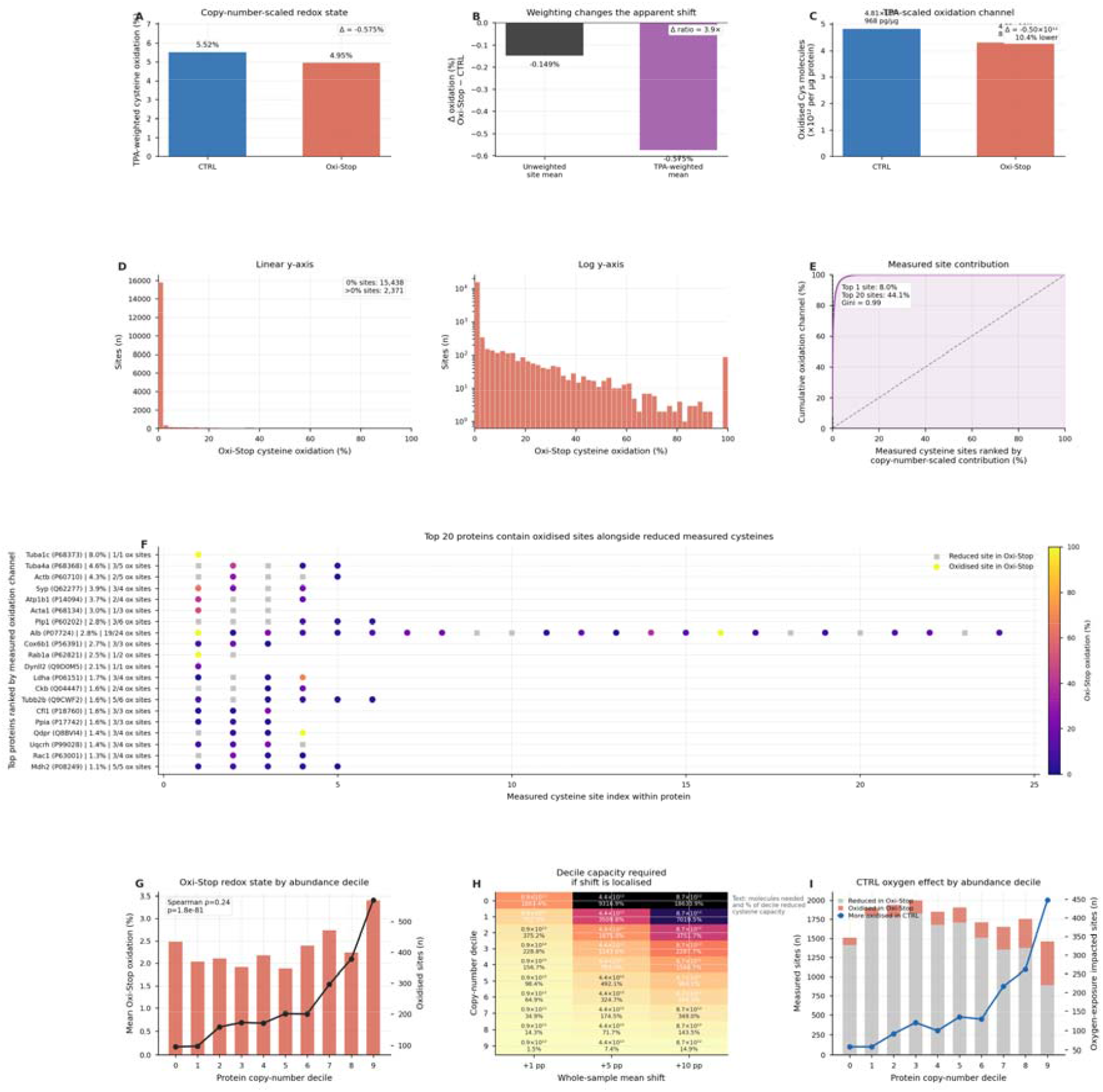
Copy-number scaling resolves cysteine oxidation as a sparse molecular redox channel. **(A)** Total protein approach (TPA)-weighted cysteine oxidation in CTRL and Oxi-Stop mouse brain hemispheres. Copy-number scaling increased the estimated cysteine redox state compared with the unweighted site mean and resolved a larger between-group difference. **(B)** Comparison of the unweighted and TPA-weighted group difference. The apparent Oxi-Stop effect increased from −0.1485 percentage points using the unweighted site mean to −0.575 percentage points after TPA-based scaling, corresponding to a 3.9-fold amplification of the estimated redox shift. **(C)** Molecular scaling of the cysteine oxidation channel. CTRL contained an estimated 4.81 × 10^12^ oxidised cysteine molecules per µg protein, compared with 4.31 × 10^12^ in Oxi-Stop, corresponding to 5.01 × 10^11^ fewer oxidised cysteine molecules per µg protein after oxygen exclusion. **(D)** Distribution of endogenous cysteine oxidation in Oxi-Stop samples shown on linear and log-scaled y-axes. Most quantified cysteine sites occupied the fully reduced boundary, whereas a smaller oxidised fraction formed a long right-skewed tail. **(E)** Ranked cumulative contribution of individual cysteine sites to the copy-number-scaled oxidation channel in Oxi-Stop. The top-ranked site accounted for 8.0% of the measured oxidation channel and the top 20 sites accounted for 44.1%, demonstrating strong molecular concentration of the oxidation signal. **(F)** Site-level architecture of the top 20 proteins contributing to the measured oxidation channel. Each row represents one protein and each point represents a measured cysteine site. Oxidised sites are shown alongside reduced sites, demonstrating that high-contribution proteins are not globally oxidised but instead carry one or a few high-contribution oxidised cysteines within an otherwise largely reduced cysteine background. **(G)** Oxi-Stop redox state by protein copy-number decile. Higher-abundance deciles contained more oxidised cysteine sites and showed higher mean cysteine oxidation, indicating that the endogenous oxidation channel is abundance structured. **(H)** Estimated decile capacity required to shift the whole-sample cysteine redox mean by 1, 5, or 10 percentage points. Values show the number of additional cysteine molecules required and the corresponding percentage of the reduced cysteine capacity within each copy-number decile. **(I)** Distribution of reduced, oxidised, and oxygen-exposure-sensitive cysteine sites across protein abundance deciles. Oxi-Stop-protected sites were detected against a much larger background of reduced cysteines, showing that the storage-induced signal acts within a sparse redox landscape rather than uniformly across the measured cysteine proteome.

We next used the Oxi-Stop condition to define the minimally distorted endogenous cysteine oxidation landscape. This landscape was highly sparse. Of 17,809 quantified cysteine sites, 15,438 were fully reduced, whereas only 2,371 showed any measurable oxidation. The distribution of Oxi-Stop oxidation was strongly right-skewed, with a large reduced boundary and a long oxidised tail (**Fig. 7D-E**). Copy-number scaling showed that this sparse site-level structure was also highly concentrated at the molecular level. The top-ranked cysteine site accounted for 8.0% of the measured oxidation channel, the top 20 sites accounted for 44.1%, and the top 20 proteins accounted for 53.9% (**Fig. 7F**). Thus, the endogenous cysteine oxidation channel was not diffusely distributed across all detected cysteines; it was concentrated into a narrow set of high-contribution sites and proteins.

Consistent with an information encoding oxidation channel, these high-contribution proteins also contained reduced sites (**Fig. 7G**). Mapping all measured cysteine sites across the top 20 contributing proteins showed that oxidised cysteines coexisted with many reduced cysteines on the same proteins. In other words, these proteins dominated the copy-number-scaled oxidation channel because one or a few sites contributed strongly, not because the entire cysteine complement of the protein was oxidised. This distinction is essential: copy-number scaling identifies where the quantitative oxidation signal resides, but the underlying proteoform space remains largely reduced [38–40].

The abundance structure of the oxidation channel was further supported by decile analysis. Higher copy-number deciles contained more oxidised sites and showed higher mean Oxi-Stop oxidation than lower-abundance deciles. The oxygen-exposure effect in CTRL also mapped onto this abundance structure, with Oxi-Stop-protected sites detected against a much larger background of reduced cysteines within the same abundance deciles. Therefore, storage-induced oxidation did not simply add a uniform offset to the cysteine proteome. It acted within an already sparse, abundance-structured redox landscape, where small changes in specific high-copy-number proteins or sites can produce large shifts in the molecular oxidation channel.

Together, these analyses show why cysteine redox proteomics cannot be interpreted from fractional occupancy alone. Percentage oxidation identifies where oxidation is detectable; copy-number scaling identifies where oxidation quantitatively resides. In mouse brain, Oxi-Stop revealed that endogenous cysteine oxidation is a sparse molecular channel dominated by a small number of high-contribution sites on abundant proteins, while the surrounding cysteine proteome remains overwhelmingly reduced. This converts cysteine oxidation from a dimensionless site-occupancy measure into a unit-scaled, abundance-weighted molecular signal.

## Discussion

Here, we demonstrated that exposure of tissue samples to atmospheric oxygen, during only 14 days of cryopreservation, induced pervasive oxidation, confounding redox proteomics. To address this problem, we developed and validated the redox capture (ReCap) workflow, comprising Oxi-Stop-enabled Oxi-DI for deep, quantitative redox proteomics with minimal exogenous oxidation. Spurious redox drift arising from exogenous oxidation during sample storage was minimised by Oxi-Stop before Oxi-DIA provided deep, copy number-scaled cysteine redox proteomics.

Oxi-DIA resolved a long-standing quantitative limitation in redox proteomics by simultaneously measuring residue-level cysteine oxidation states and protein copy numbers, without requiring additional sample enrichment steps. This enabled, for the first time, the relative copy number scaling of cysteine redox states. Just 20 cysteines hosted by abundant proteins in the mouse brain proteome accounted for ∼44% of all oxidised cysteine molecules in mouse brain, with tubulin alpha-1C chain Cys295 alone contributing 8% of the total cysteine oxidation. Hence cysteine oxidation is concentrated more narrowly in a subset of the proteome than previously appreciated.

The copy-number-weighted quantification of protein redox values is a prerequisite for biologically meaningful interpretation of the redox proteome data. For example, even if a large, 20% increase in the oxidation level occurred in a thousand different low abundance proteins, it would barely alter the mean global redox state of the tissue proteome. Hence, a major redistribution of the oxidation channel (i.e., numbers of oxidised proteins), is possible without necessarily resulting in “oxidative stress”, as defined by an increase in the mean cysteine oxidation of the proteome.

The demonstration here that a major preanalytical variable (i.e., storage-induced oxidation) that confounded cysteine redox proteomics has immediate practical consequences. The data show that atmospheric oxygen exposure, during even short periods of cryopreservation, amplifies cysteine oxidation at biologically important sites. This exposure spuriously amplified the difference in the global cysteine redox state between the left and right hemisphere by one order of magnitude. Hence, retrospective reinterpretation of conclusions from previous redox proteomics datasets, derived from stored tissues that lacked this control, may be warranted.

To conclude, we recommend that Oxi-Stop is widely adopted for tissue storage and used to minimise atmospheric oxygen exposure-induced redox drift during cryopreservation in biobanks. This simple and inexpensive strategy is expected to materially improve the interpretability of future redox proteomic studies. By enabling deep, copy-number scaled cysteine redox proteomics with minimal exogenous oxidation, we anticipate that the ReCap workflow will lead to advances in current understanding of redox biology.

## Methods

### Experimental models

#### Mouse model

All procedures were performed in accordance with the Animals (Scientific Procedures) Act, 1986 and approved by the University College London Ethical Review Process Committee. Mouse husbandry and health monitoring were as previously described [41]. Male wild-type mice were generated by backcrossing to C57BL/6J (Charles River, UK, RRID:IMSR_JAX:000664). Mice were culled at ten weeks of age by a schedule 1 procedure, brain hemispheres were rapidly dissected and then either placed in a cryovial (CTRL) or tightly wrapped in layers of aluminium foil (Oxi-Stop) before placement in a cryovial, and snap frozen in liquid nitrogen. The samples were stored at −70℃ for 14 days and then processed identically for Oxi-DIA altogether (i.e., to exclude batch effects). Per previous work [12,42], tissues were mechanically lysed in buffer using beads (Percellys, 6000 *g* for 30 s at 4°C).

In this design, the left and right hemisphere corresponded to CTRL and Oxi-Stop, respectively. In a separate experimens, left and right hemispheres from male mice both wrapped in aluminium foil (Oxi-Stop) were analysed by Oxi-DIA.

#### Cell model

HEK293 cells (RRID:CVCL_0045) previously described [43] were purchased from ATCC and cultured in Dulbecco’s modified Eagle’s medium supplemented with 10% fetal bovine serum (ThermoFisher, UK, #A5209402), 2 mM glutamine (ThermoFisher, UK, #25030149) and 100 µg/ml penicillin-streptomycin (ThermoFisher, UK, #15070063). Cells were trypsinised in TrypleE (ThermoFisher, UK, #12604013) following DPBS washes and plated in appropriate cell culture plate. Cells were routinely tested for mycoplasma (MycoStrip, InvivoGen, UK). Approximately 2x10^6^ cells/ml were harvested per biological replicate and then lysed immediately. These cells were lysed by passing them 10-12 times through a 21G needle.

Regardless of the model or the assay, the samples were processed in 2 ml Protein LoBind Eppendorf tubes to avoid passive protein loss (VWR, UK, #022431102).

### DTNB assay

To determine if aluminium foil donated electrons to a model thiol, we used DTNB (Sigma, UK, #D8130). If DTNB is reduced, then measurable product 2-nitro-5-thiobenzoic acid (TNB) is produced. For the aqueous assay, a strip of aluminium foil was immersed in reaction buffer (10 mM DTNB, 0.1 sodium phosphate, pH 8, 1 mM EDTA) inside a 2 ml Eppendorf for 30-min at room temperature in the dark. As a negative control, the reaction buffer was incubated for 30-min in just the Eppendorf. Hence, plastic served as a passivated negative control material. As a positive control, the reaction buffer was incubated with 5mM TCEP for 30-min. For the solid-state assay 4 mg of DTNB was either wrapped in aluminium foil or stored in an Eppendorf (negative plastic control) for 30-min at room temperature in the dark. Thereafter, the DTNB was hydrated in reaction buffer for measurement. For both the aqueous and solid-state assays, the production of TNB was measured in a microplate reader at 412 nm. After subtracting the background signal at 412 nm (derived from just reaction buffer), the signal of TNB was calculated in arbitrary units (AU).

### Fluorescent thiol assay

To test approaches for preventing lysis-induced cysteine oxidation, HEK293 cells were lysed in 2% ultra-pure SDS and 100 mM HEPES (pH 7.4) in the presence or absence of metal ion chelators. EDTA (2 mM, Sigma, UK, #03690), DTPA (1 mM, Sigma, UK, #D6518) and neocuprine (0.1 mM, Sigma, UK, #121908) were added individually and collectively to the lysis buffer, and samples were incubated for 30 min at 37 °C.

Cysteine oxidation was quantified as a loss of AlexaFluor™-488-C_5_-maleimide (F-MAL, ThermoFisher, UK, #A10254) labelling [44], which labels reduced cysteines. Samples were alkylated with 5 mM F-MAL immediately after 30-min, and the excess reagent was removed by using spin-column desalting [45]. Fluorescence was measured by using a microplate reader and normalised to total protein (colorimetric detergent-compatible Bradford assay). Cysteine labelling was calculated as:

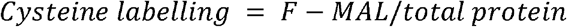

To assess the contribution of metal-dependent oxidative reactions during lysis, parallel samples containing chelators were processed with or without immediate addition of F-MAL. The assay was also used to quantify oxidation induced by two commonly used lysis accelerants: sonication (10 x 30 seconds) and boiling (5 min at 90 °C). Boiling could induce the homolysis of disulfide bonds, producing sulfur radical species [28]. Meanwhile acoustic cavitation during sonication can produce the hydroxyl radical [27]. All reactions were performed in the dark to minimise light-induced side-reactions, such as superoxide anion production via the excited flavin species or the photoexcitation of the fluorophore [46,47].

### Orthogonal benchmarking

The Oxi-DIA paired with DIA-NN workflow described below was orthogonally benchmarked in HEK293 cells. HEK293 cells lysed in NEM_L were cleaned up (see **Cysteine labelling**) and resuspended in 80 µg of TCEP-immobilised magnetic beads (BioClone, USA, #BH101) for 1 h at 40°C, the samples were then magnetised to remove the beads. The sample was then transferred to a fresh tube containing 3 mg of 2-pyridyldithiol-derivatised magnetic beads (BioClone, USA, #FX104) and incubated for 3 h at 37°C. The 2-pyridyldithiol group selectively and covalently enriches proteins with at least one newly reduced oxidised cysteine residue via a disulfide bond [48]. The beads were magnetised and the eluent was retained as the reduced fraction. Bound proteins were released from the solid support enriched (oxidised) fraction using TCEP. After digesting the fractions, their peptide content was determined (see **Cysteine labelling**).

In separate experiments, F-MAL was added to the eluent in the presence of TCEP (see **Fluorescent thiol assay**). The lack of F-MAL labelling demonstrated that the eluent was reduced (pre-labelled with NEM_L) and devoid of residues that could be reduced and then labelled with the maleimide-derivatised fluorophore. Reciprocally, F-MAL was added to the enriched fraction in the presence of TCEP. Before the fluorescent thiol assay was implemented, excess fluorophore was removed (see **Cysteine labelling**).

To quantitatively benchmark percentage redox states, we compared the Oxi-DIA computed global redox state of HEK293 cells against an orthogonal global-mode RedoxiFluor assay [29,49]. In these experiments, Oxi-DIA isotopically encoded the redox state via the NEM_L and NEM_H labels. In contrast, RedoxiFluor spectrally encoded the redox state via fluorescent AlexaFluor™488-C_5_-maleimide (reduced, F-MAL1) and AlexaFluor™647-C_2_-maleimide (F-MAL2, oxidised, ThermoFisher, UK, #A20347) labels. The redox state of the cysteine proteome was determined by measuring the F-MAL1 (494-518 nm) and F-MAL2 (651-671) fluorescence in a microplate reader. After blank subtraction, the F-MAL1 and F-MAL2 signals (*v*) were corrected via:

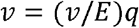

where *E* and *q* are the excitation coefficient and quantum yield, respectively. The percentage cysteine oxidation was computed as

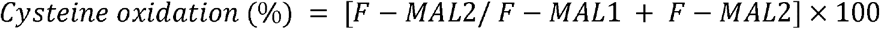

### Cysteine labelling

To label cysteines for Oxi-DIA, samples were lysed in fresh light-protected lysis buffer (2%SDS [ThermoFisher, UK, #15553027), 100 mM HEPES (pH 7.4[Sigma, UK, #0000338713), 2 mM EDTA, 1 mM DPTA, and 0.1 mM neocuprine) supplemented with 25 mM light *N*-ethylmaleimide (NEM_L, Sigma, UK, #04260). The NEN_L was added just before use to prevent aqueous hydrolysis to cysteine unreactive maleimic acid [50]. After clarifying the lysate by centrifugation (5,000 *g* for 2 min), the samples were incubated for 1 h in the dark at 37°C to enable NEM-L to alkylate reduced cysteines via a thioether bond.

After completing the NEM_L labelling reaction, the sample was cleaned up via an organic solvent-based extraction procedure [51]. The pellets were subsequently resuspended in lysis buffer supplemented with 10 mM of neutral-TCEP (ThermoFisher, UK, #77720) to reduce reversibly oxidised cysteines via nucleophilic phosphine chemistry [52]. After an incubation period of 20 min, 15 mM heavy *N*-ethyl-d_5_-maleimide (NEM_H, Sigma, UK, #692905) was added to the lysate to alkylate newly reduced (oxidised) cysteines [53]. After completing the differential alkylation reactions, the samples were cleaned via organic phase extraction.

The pellets were resuspended in 50 mM ammonium bicarbonate and digested overnight at 37°C in the dark using 1-10 µg sequence-grade Trypsin and LysC (ThermoFisher, UK, #A40009). The digestion reaction was quenched using 1% FA. The volatile ABC salt was removed via drying the peptides at 30°C to dryness in a speedvac. Peptides were resuspended in 1% DMSO and 0.1%FA. Peptide concentration was determined using a fluorescent microplate-based assay (ThermoFisher, UK, #23290).

### Oxi-DIA standard curve

To prepare the redox standard curve, mouse brains were lysed in the presence of 15 mM TCEP and then cleaned up (see **Cysteine labelling**). The resultant pellets were resuspended in 5 mM TCEP and either labelled with 50 mM NEM_L (reduced standard) or 10 mM NEM_H (oxidised standard). The two standards were then cleaned up, digested, and the resultant peptides were desalted then resuspended (0.1% formic acid [FA] & 1% dimethylsufoxide [DMSO]). As the peptide concentration exceeded the quantitative limit of the peptide assay, the standards were prepared as follows 0 (10 µl NEM_H), 20 (8 µl NEM_H + 2 µl NEM_L), 40 (6 µl NEM_H, 4 µl NEM_L), 60 (4 µl NEM_H, 6 µl NEM_L), 80 (2 µl NEM_H, 8 µl NEM_L), 90 (1 µl NEM_H, 9 µl NEM_L), 95 (0.5 µl NEM_H, 9.5 µl NEM_L), and 100 (10 µl NEM_L). Each standard was then diluted in 9 µl loading buffer (0.1%FA, 1%DMSO) and supplemented with 1 µl of indexed retention time (iRT, Brucker Daltonics, #1816351) standard peptide mix.

### Oxi-DIA

To measure the cysteine redox state via Oxi-DIA, 500 nanogram of peptides were loaded onto a Vanquish Neo U-HPLC coupled to an Orbitrap Astral™ mass spectrometer[22,26], which was mass calibrated in positive ion mode before each run. After passing through a 5 cm trap column, the samples were loaded onto a 25 cm IonOptics C_18_ column and resolved over 42-min with the ion source set to between 1600-2500 V. Mobile phase A and B were 0.1%-FA and 80% acetonitrile (ACN) in 0.1%-FA, respectively. By adjusting the mobile phase, the peptides were resolved using a nonlinear gradient 5-80% ACN. MS^1^ (intact peptides) spectra were acquired in the Orbitrap every 0.6 seconds at the full instrument resolving power of 240,000. After fragmenting peptides via high-energy collisional induced dissociation at 25%-energy, MS^2^ spectra were collected in the Astral across the 380-980 m/z range at full instrument resolving power, automatic gain control of 500%, and a maximal injection time of 3 milliseconds. The 380-980 m/z range was split into 300, 2-Thompson windows for narrow DIA [54].

### DIA-NN

Thermo raw files were analysed in DIA-NN (v.2.2) [25] using an *in silico* predicted spectral library generated from a species-specific FASTA file. The spectral library was generated with the following instructions inputted into the command line interface: “-*-var-mod NEM_L,125*.*047,C*”, “*--var-mod NEM_H, 130*.*076,C*”, and “*--original-mods*”. In addition, variable N-terminal methionine excision and up to 1 missed tryptic digestion were allowed. The maximum number of variable modifications per peptide was set to 3 and the peptide length range considered was 5-35 amino acids. For the quantitative search, the raw files were searched against the *in silico* generated spectral library. For this search, “*--export quant*” was added to the command line interface. The other search settings remained the same. Match between runs (MBR) and peptidoform scoring mode were enabled. In addition, quant-UMS (precision) was selected to minimise ratio distortion [55]. Per recent work [13], the peptide and protein false discovery rate (FDR) was set to 0.01 and sites with localisation probabilities of 0.9 were computationally analysed.

### Computational analysis

DIA-NN output files were processed using custom Python scripts implemented in Jupyter notebooks and executed within a high RAM Google Colab runtime [40]. The scripts and their relevant outputs (e.g., data files) are available at https://github.com/JamesCobley/ReCap.

#### Validation of Oxi-DIA

Only cysteine-containing peptides were retained for analysis. Peptides were deduplicated per LC–MS run and stripped sequence to prevent repeated quantification of identical peptide entries within the same acquisition. Modified sequences were parsed to assign isotopic label state (NEM_L or NEM_H). For each run, total cysteine-containing peptides, light-labelled peptides, heavy-labelled peptides, and unlabelled peptides were enumerated. Observed fractional reduction was computed from the ratio of light-labelled peptide counts to total labelled peptides (MS^2^-level proxy) and to total cysteine peptides including unlabelled entries (MS^1^-level proxy). A combined estimate was calculated as the arithmetic mean of MS^1^ and MS^2^ proxies. Expected fractional reduction values for the mixing series were derived from nominal mixing percentages, adjusted for empirically estimated unlabelled peptide fractions obtained from pure 0% and 100% reference samples. Linear interpolation was used to model the expected unlabelled contribution across intermediate mixing states. Quantitative performance metrics including R^2^, RMSE, MAE, MAPE, residual bias, and residual standard deviation were computed using scikit-learn. Residuals were defined as observed minus expected cysteine redox state.

#### Pair-matched mouse-brain analysis

Peptide entries were parsed from the Protein.Sites field to extract protein accession, residue identity, and residue position, and only cysteine-containing sites were retained. Fragment-level quantitative columns (Fr.X.Quantity) were summed within each peptide entry to generate a total MS^2^ signal. Modified peptide sequences were then classified according to isotopic label state as light (NEM_L) or heavy (NEM_H). For each run and cysteine site, MS^2^ signal was summed separately for light- and heavy-labelled species and cysteine oxidation was calculated as:

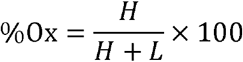

where *H* and *L* denote the total heavy- and light-labelled MS^2^ signal, respectively. Sites shared across all included biological replicates were retained for downstream comparison. Mean site-level oxidation values for each condition were calculated as the arithmetic mean across biological replicates:

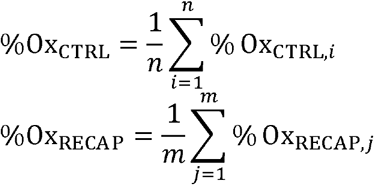

and the site-level redox difference between conditions was defined as:

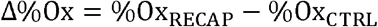

A site-level output table containing per-site mean oxidation values together with biological replicate values was exported for downstream identity, boundary-state, and annotation analyses. The same computational process was used to analyse the cysteine redox proteomic data from the HEK293 cell model.

Sample-level weighted mean cysteine redox state was calculated on the common shared cysteine set to ensure comparability across samples. For each shared cysteine site, the Oxi-DIA percent oxidation value was assigned to the MaxLFQ-derived abundance of its parent protein group in the same run. Protein abundances were collapsed to one value per run and protein group by retaining the maximum repeated protein-level quantity where necessary. Weighted mean redox was then computed as

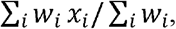

with *x*_*i*_ denoting site-level percent oxidation and *w*_*i*_ the corresponding protein-group abundance. This yielded one weighted mean redox value per sample, which was used for group-wise comparison

#### Resources

We contextualised the measured cysteine redox sites against the broader mouse cysteine landscape by generating reference resources from the canonical Mus musculus UniProt proteome. First, the canonical mouse FASTA was parsed using Biopython to enumerate proteins, protein length, and cysteine content for each sequence. For each protein, the total number of cysteine residues was counted directly from the primary amino acid sequence, allowing calculation of the total number of proteins represented, the number containing at least one cysteine residue, and the total cysteine content of the canonical proteome. Second, UniProtKB JSON records were retrieved for each protein accession using the UniProt REST API. Positional feature annotations were parsed and mapped onto cysteine coordinates within each protein sequence. Overlap between cysteine positions and annotated feature intervals was used to identify cysteines corresponding to selected functional categories, including active sites, binding sites, metal-binding sites, disulfide bonds, and zinc-finger regions. Annotation counts were aggregated at both the protein level and the global proteome level. Third, a site-level cysteine functional resource was generated by extracting positional annotations for selected UniProt feature classes and retaining only annotated cysteine residues. For each annotated cysteine site, accession, gene symbol, protein name, organism, review status, cysteine position, feature type, feature span, ligand metadata, and evidence fields were recorded where available.

A protein-level annotation resource was generated from the cysteine summary table. Proteins containing at least one cysteine residue were retained and annotated using UniProtKB JSON records retrieved through the UniProt REST API. For each protein, accession, primary gene name, recommended protein name, organism, and review status were extracted. UniProt cross-references were parsed to collect Gene Ontology (GO), KEGG, and STRING annotations. GO terms were classified into Cellular Component, Biological Process, and Molecular Function categories based on UniProt GO term prefixes. A protein-level master table was generated containing cysteine content, counts of annotated functional cysteines, and the number of associated GO, KEGG, and STRING annotations. A companion long-format annotation table was also produced containing one row per protein-annotation pair.

#### Site-level analysis

To derive site-level insights, open-sourced scripts from the Oxi-Shapes resource [35] were adapted. A symmetry analysis was performed on shared cysteine sites using mean site-level oxidation values for CTRL and RECAP. For each site *i*, the redox difference was defined as

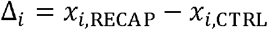

where *x*_*i*,CTRL_and *x*_*i*,REDCAP_ denote the mean percentage oxidation across biological replicates in each condition. Sites were classified as identical when Δ_*i*_ =0 and nonidentical when Δ_*i*_ .≠ 0 . Identity analysis was performed without tolerance; sites were considered identical only when the mean oxidation values were numerically equal between conditions. Identical sites were further partitioned into shared 0% states, shared 100% states, and shared partial states. Nonidentical sites were partitioned by direction (Δ_*i*_ >0 or Δ_*i*_ < 0) and by state class. A nonidentical site was considered switch-like if either condition lay at a boundary value (0% or 100%), and toggle-like if both values lay strictly within the partial interval 0 < *x* < 100. The same framework was also applied to the subset of sites exhibiting nonzero oxidation in either or both conditions.

Site-level redox classifications were overlapped with a curated UniProt-derived cysteine functional annotation resource by matching protein accession and cysteine position. This produced a merged table containing all quantified sites together with functional annotations where present. Functionally annotated nonidentical sites were defined as nonidentical sites with at least one matched annotation and were summarised by direction of change, feature type, and gene identity.

Shared quantified cysteine sites were first collapsed to the protein level by accession. A protein was classified as nonidentical if any constituent site satisfied NonIdentity = TRUE, and as identical otherwise. Gene Ontology (GO) annotations were then mapped to measured proteins using the UniProt-derived protein annotation resource, with duplicate protein-term pairs removed. For each GO term, enrichment in the nonidentical protein set relative to the identical protein set was assessed using a two-sided Fisher’s exact test on the contingency table

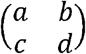

where *a* and *c* denote the numbers of nonidentical and identical proteins annotated with the term, respectively, and *b* and *d* denote the corresponding numbers without the term. Resulting *p*-values were corrected for multiple testing by the Benjamini–Hochberg method. For significant GO terms, the associated site-level redox dataset was then summarised by the number and proportion of nonidentical sites, direction of Δ%Ox, and mean, median, and absolute mean redox differences across sites.

### Statistical analysis

The statistical test used to analyse a dataset is reported in the figure legends. These tests were implemented in a Google Colab or GraphPad Prism environment as appropriate. In all cases (including FDR corrections), alpha was set to < 0.05.

## Data availability

The raw MS files will be deposited to PRIDE. The outputs resulting from our computational analysis are available online, such as the oxidation matrix table reporting all site level data (i.e., https://github.com/JamesCobley/ReCap/blob/main/Oxi_Stop/shared_cysteine_redox_matrix_CTRL_vs_REDCAP.csv).

