## Supplemental Information for "ReCap enables deep, copy number-scaled cysteine redox proteomics with minimal exogenous oxidation"

**Supplementary figures**

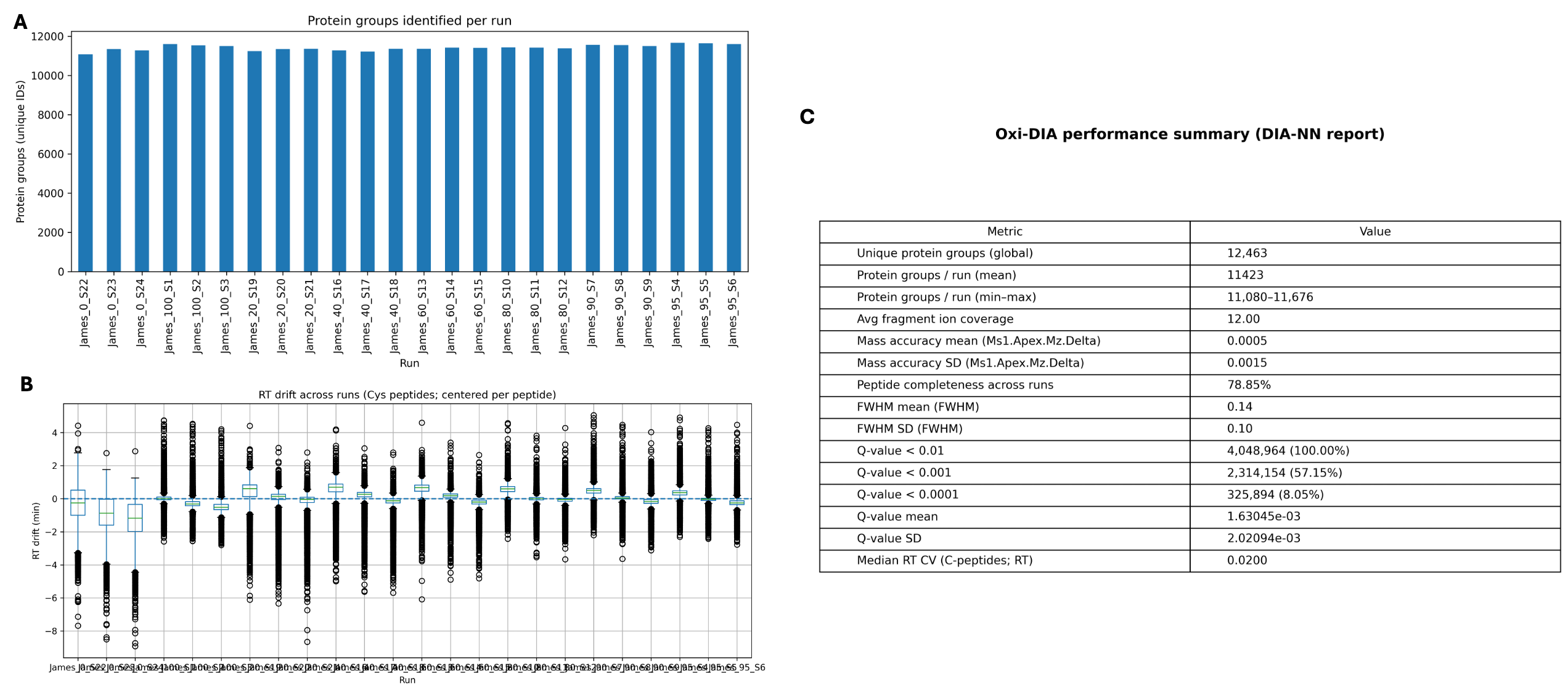

**Supplementary Figure 1 | Analytical performance and chromatographic stability of Oxi-DIA.**
(A) Number of unique protein groups identified per run from DIA-NN analysis of the Oxi-DIA acquisition series, demonstrating consistent proteome coverage across runs.
(B) Retention time (RT) stability assessed using cysteine-containing peptides. For each peptide, RT values were centred by subtracting the median RT across runs, and distributions of centred RT values are shown per run. The median RT coefficient of variation (CV) across cysteine-containing peptides was 0.020, indicating high chromatographic reproducibility.
(C) Summary of global performance metrics derived from the DIA-NN report, including unique protein groups identified, fragment ion coverage, mass accuracy, peptide completeness across runs, peak width (FWHM), and q-value distributions. All metrics were computed directly from the DIA-NN output using a reproducible analysis pipeline (<https://github.com/JamesCobley/ReCap/blob/main/Oxi-Dia/suppfig1_qc_pipeline.py>).

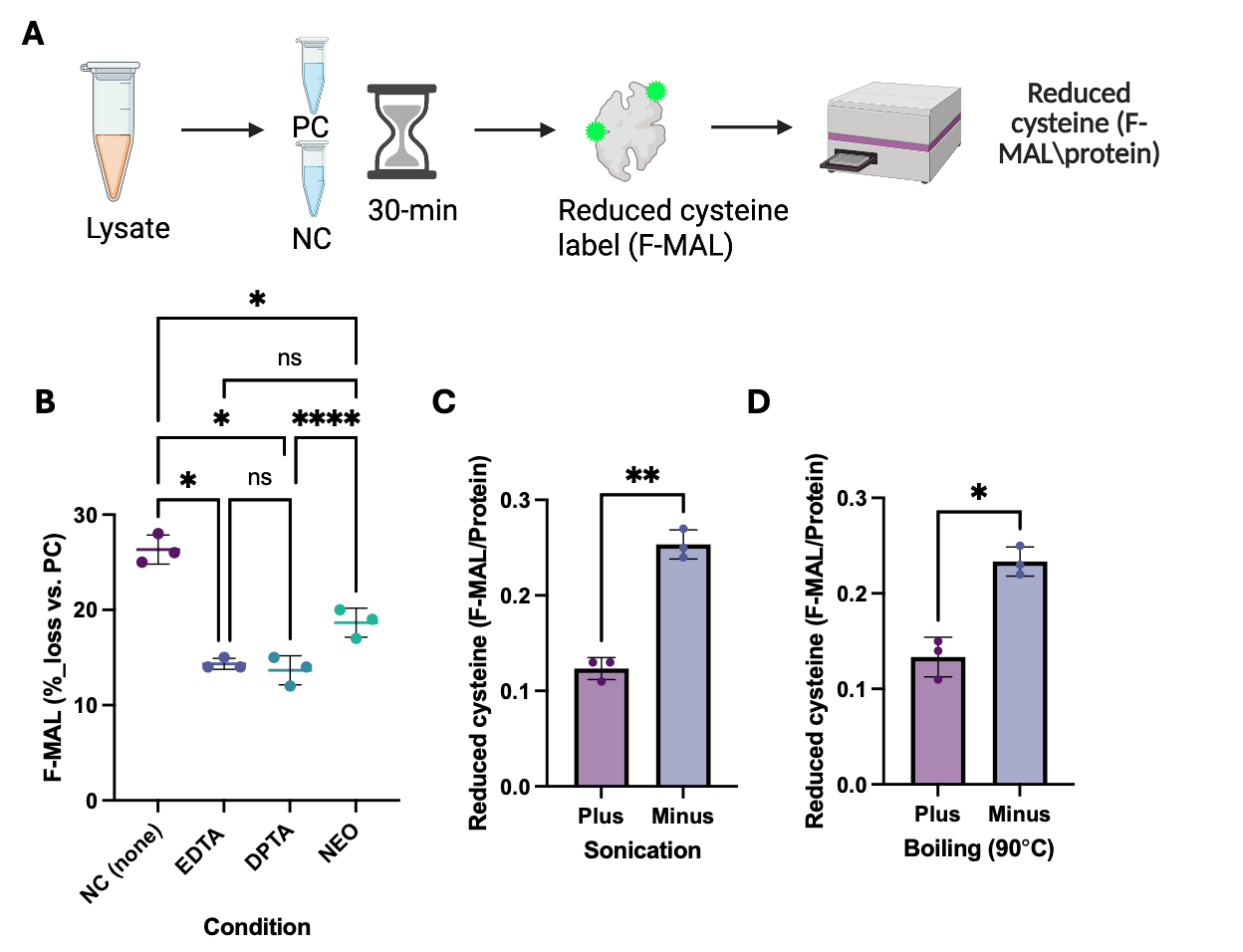

**Supplementary Figure 2 | Extended validation of Oxi-DIA.**
(A) Schematic illustration of the fluorescent thiol assay used to determine the effectiveness of strategies to minimise lysis-induced cysteine oxidation. Lysis-induced cysteine oxidation is measured as a loss of fluorescent maleimide (F-MAL) labelling compared to the positive control, application of the F-MAL label at the point of lysis in the presence of the chelators (EDTA, DPTA, and neocuprine).
(B) The effect of complete omission of all chelators (negative control) compared to the addition of chelator relative to the positive control (all chelators present). Statistical significance was assessed via a one-way ANOVA with FDR corrected Tukey post-hoc testing.
(C) The effect of sonication and (D) boiling on F-MAL labelling in HEK293 cell lysates.

Panels B-D report the results from *n* = 3 HEK293 cell lysates per condition. Panels E-F are *n* = 6 HEK293 lysates analysed by Oxi-DIA. Note, ns, *,**, and **** denote non-significant, $P=<0.05, P= <0.01,P=<0.0001$, respectively.

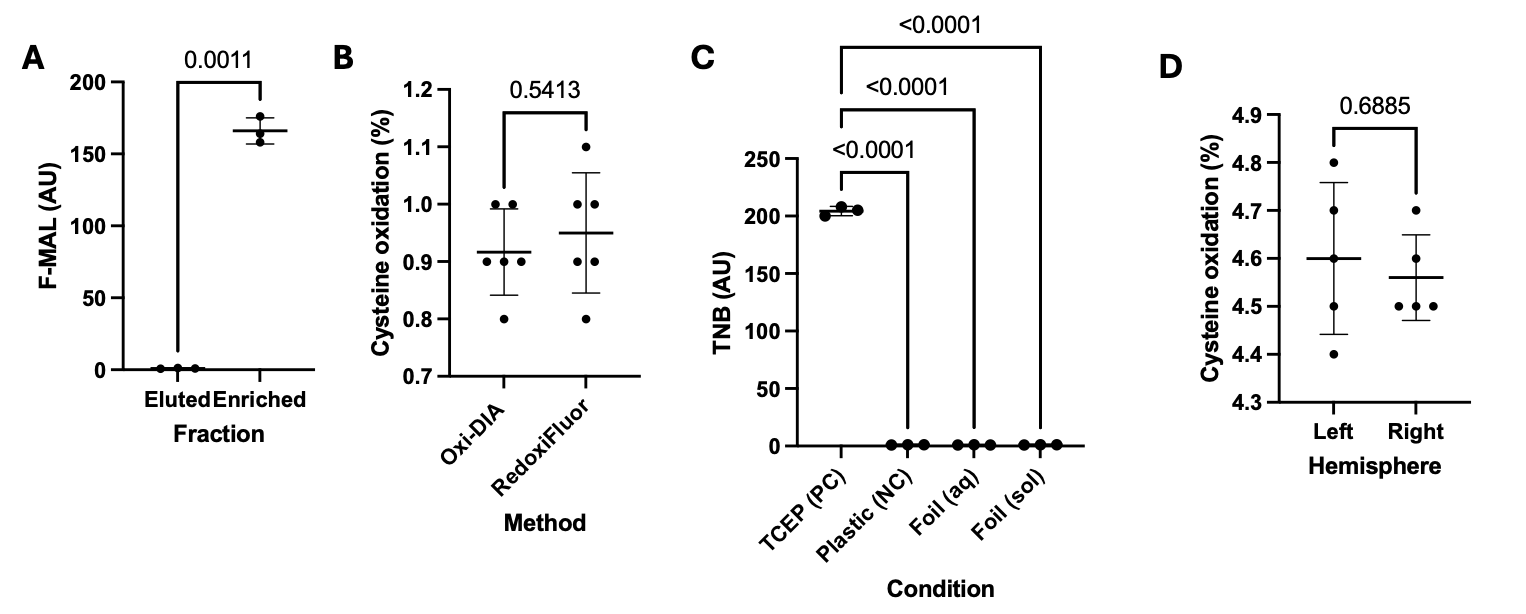

**Supplementary Figure 3 | Validation of Oxi-DIA, Oxi-Stop, and the mouse brain model.**

(A) Fluorescent maleimide (F-MAL) signal (arbitrary units, AU) in each fraction (*n* = 3). Paired t-test.

(B) The global cysteine proteome redox state in Oxi-DIA compared to RedoxiFluor in HEK293 cells (*n* = 6). Unpaired t-test.

(C) The reduction of DTNB (a model oxidised thiol) to TNB (arbitrary units, AU) by TCEP (positive control) compared to plastic (negative control) and aluminium foil under aqueous (aq) and solid state (sol). There were three biological replicates in each group. Statistical significance was assessed using a one-way ANOVA, corrected for multiple comparisons using the Dunnet method**.**

(D) No statistically significant difference (paired *t*-test) in the mean cysteine redox state in left (*n* = 5) and right (*n* = 5) mouse brain hemisphere that were both stored under the Oxi-Stop condition.

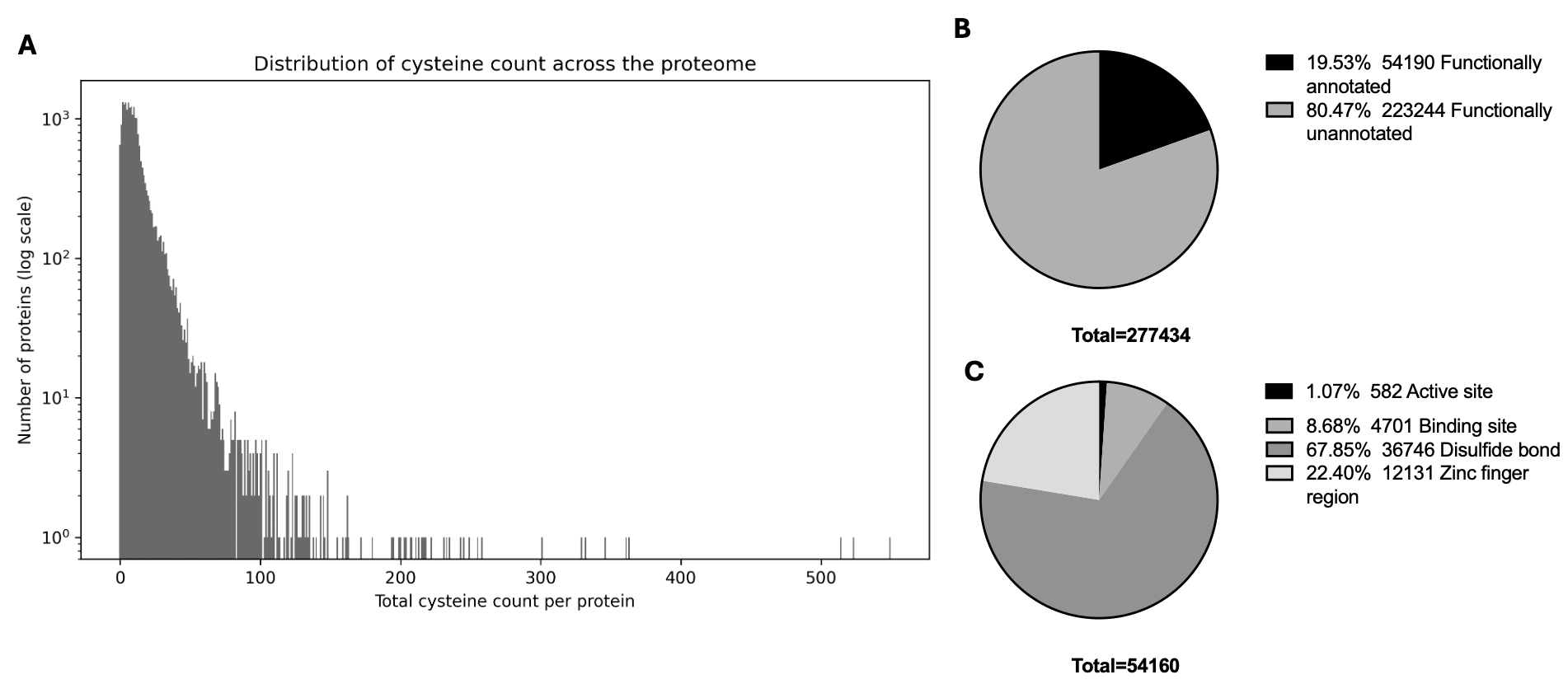

**Supplementary Figure 4 | Cysteine distribution and functionality in the mouse reference proteome.**
(A) Histogram plot of the number of protein groups by cysteine residue number in the mouse reference proteome.

(B) Pie chart of functionally annotated and unannotated cysteine residues in the reference mouse proteome based on a computational search of the UniProt database per the methods.

(C) Pie chart of the distribution of annotated cysteine residues by function class
